# Insulin Signaling-independent Activation of DAF-16 Shapes the Transcriptome during Normal Aging

**DOI:** 10.1101/284083

**Authors:** Shang-Tong Li, Han-Qing Zhao, Pan Zhang, Chung-Yi Liang, Yan-Ping Zhang, Ao-Lin Hsu, Meng-Qiu Dong

## Abstract

The roles and regulatory mechanisms of transriptome changes during aging are unclear. It has been proposed that the transcriptome suffers decay during aging owing to age-associated down-regulation of transcription factors. In this study, we characterized the role of a transcription factor DAF-16, which is a highly conserved lifespan regulator, in the normal aging process of *Caenorhabditis elegans*. We found that DAF-16 translocates into the nucleus in aged wild-type worms and activates the expression of hundreds of genes in response to age-associated cellular stress. Most of the age-dependent DAF-16 targets are different from the canonical DAF-16 targets downstream of insulin signaling, indicating that activation of DAF-16 during aging is not due to reduced insulin signaling from DAF-2. Further analysis showed that it is due to the loss of proteostasis during aging, at least in part. We also found that without *daf-16*, dramatic gene expression changes occur as early as on adult day 2, indicating that DAF-16 acts to stabilize the transcriptome during normal aging. Our results thus reveal that normal aging is not simply a process in which the gene expression program descends into chaos due to loss of regulatory activities; rather, there is active transcriptional regulation that fights aging.

## Introduction

Aging is characterized by a gradual deterioration of physiological functions and an increase of the probability of death. Old age is the single most important risk factor for all gerontological diseases from arthritis to Alzheimer’s. Undoubtedly, aging is a process accompanied by numerous biological changes at different levels, from the molecular to the systemic and organismal. However, the aging process has not been characterized comprehensively and quantitatively. As such, for practical reasons, aging is measured not by a metric of the process but by its end point—death. Surely, lifespan assay is a convenient way to assess the average rate of aging, but it does not inform anything about the dynamics. For example, whether aging proceeds at an even pace or transitions through distinct phases with acceleration or deceleration is not known.

The nematode *C. elegans* has long established itself as an ideal model for aging research. Key genetic pathways that regulate aging have been discovered originally in *C. elegans* and later confirmed in other organisms including mammals, and one of which is the Insulin/Insulin-like Growth Factor 1 (IGF-1) signaling pathway, or IIS (Kenyon, Chang, Gensch, Rudner, & Tabtiang, 1993). In the *C. elegans* IIS pathway, the sole insulin/IGF-1 receptor DAF-2 negatively regulates the FOXO transcription factor DAF-16 through a cascade of kinases composed of PI3K/AGE-1, PDK/PDK-1 and AKTs. Phosphorylation by AKTs prevents DAF-16 from entering the nucleus to regulate downstream target genes. Loss-of-function mutations of *daf-2* or the downstream kinases extend lifespan in a *daf-16* dependent manner (Henderson & Johnson, 2001; Kimura, Tissenbaum, Liu, & Ruvkun, 1997; Lee, Hench, & Ruvkun, 2001; Lin, Dorman, Rodan, & Kenyon, 1997; Morris, Tissenbaum, & Ruvkun, 1996; Ogg et al., 1997; Paradis, Ailion, Toker, Thomas, & Ruvkun, 1999)

The *C. elegans* aging process has been characterized to some extent, including behavioral declines at the organismal level, gross morphological changes (e.g. length, texture) (Pincus, Smith-Vikos, & Slack, 2011), fine structural changes (Herndon et al., 2002; McGee et al., 2011; Regmi, Rolland, & Conradt, 2014) and molecular changes (Budovskaya et al., 2008; Golden & Melov, 2004; Lund et al., 2002; Walther et al., 2015). Among the molecular changes that are fundamental to other types of changes occurring in the aging process, gene expression changes are relatively easy to measure, thanks to the microarray technology in the past and deep sequencing at present. Previously, by analyzing transcriptome of *C. elegans* on adult days 1, 3, 5, and 10 at 20 °C, Rangaraju *et al*. identified transcriptional drift, which refers to loss of stoichiometry between mRNAs of the same functional group and loss of youth-associated co-expression patterns in aging worms, as an age-associated degenerative process. Slowing down aging by knocking down *daf-2* restricted the progression of transcriptional drift, arguing that transcriptional drift is closely related to aging (Rangaraju et al., 2015). However, it is not known how IIS contributes to the normal aging process. A recent study has found that the amount DAF-2 protein increases from adult day 1 through day 10 (Tawo et al., 2017), but it is still unclear whether there is a decrease of transcriptional activity of *daf-16* in the normal aging process, and if there is, how much it contributes to transcriptional drift in aging *C. elegans*.

A few studies have suggested a possible link between transcriptome deterioration and age-associated inactivation of transcription factors. One study showed that deactivation of a GATA factor *elt-2* is responsible for age-associated down-regulation of more than 200 genes in wild-type (WT) worms assayed on adult days 2, 5, 8, and 11 at 25 °C (Mann, Van Nostrand, Friedland, Liu, & Kim, 2016). *elt-2* plays an essential role in the development of the intestine, indicating that it is selected under strong selective pressure during evolution. Loss of the *elt-2* transcriptional network after *C. elegans* has reached adulthood is consistent with the idea that transcriptional drift plays a role in age-associated physiological deterioration. In comparison to that of *elt-2*, age-dependent repression of *hsf-1* is strikingly early—within the first 12 hours of adulthood at 20 °C, which is closely connected to the collapse of proteostasis during aging (Labbadia & Morimoto, 2015).

In this study, prompted by the observation of nuclear accumulation of DAF-16 in mid-aged *C. elegans* adults, we carried out mRNA-seq analysis of WT and *daf-16 (null)* mutant worms every 24 hours from adult day 1 through day 7. Specifically, we set out to address three questions. First, whether DAF-16 is activated during aging, and when? Second, what activates DAF-16 during aging–is it reduction of IIS or else? Third, what is the role of DAF-16 during normal aging? Through the mRNA-seq and follow-up experiments, we verified that DAF-16 is indeed activated during the aging process of WT *C. elegans*, most prominently on adult day 6 and day 7 at 25 °C. Remarkably, DAF-16 is activated not by reduction of IIS, but in response to a loss of proteostasis and possibly other types of age-associated cellular stress. Lastly, we find that DAF-16 acts as a “capacitor” to resist age-induced perturbation of the gene expression program in the normal aging process.

## Results

### DAF-16 is activated in the normal aging process

While observing WT worms expressing a DAF-16::GFP fusion protein under standard, non-stressful conditions (20 °C, well-fed), we noticed nuclear accumulation of DAF-16::GFP in older animals but not in the younger ones (Fig. 1A and S1). Quantification of the localization patterns of DAF-16::GFP verified this observation (Fig. 1B). At 20 °C, DAF-16::GFP translocated into the nucleus starting from adult day 4. As nuclear localization typically represents DAF-16 activation (Lee et al., 2001), this result suggests that DAF-16 is activated during the normal aging process. Supporting this idea, four out of five known DAF-16 target genes showed higher mRNA levels in day-3 and day-5 WT adults than in day-1 adults, and this was dependent on *daf-16* (Fig. 1C). As a poikilothermic animal, *C. elegans* ages faster at higher temperatures. If DAF-16 activation is driven by aging, then one would predict accelerated (or deaccelerated) DAF-16::GFP nuclear accumulation in worms grown at 25 °C (or 15 °C) compared to those at 20 °C. This is indeed what we observed (Fig. 1D & 1E). A previous study had compared the transcriptomes of day-3 WT and day-3 *daf-16(null)* adult worms cultured at 15 °C or 25 °C by microarray analysis (B. Zhang et al., 2015). Based on our microscopy analysis results (Fig. 1E), we estimated that on adult day 3 DAF-16 has accumulated substantially in the nuclei of WT animals grown at 25 °C, but not in those at 15 °C. So, DAF-16 is predicted to be activated in the former, but not in the latter. Agreeing with this prediction, re-analysis of the microarray data set (B. Zhang et al., 2015) identified only 37 differentially expressed genes (DEGs) between WT and *daf-16(null)* on day 3 at 15 °C, in contrast to 316 DEGs at 25 °C.

**Figure 1.**
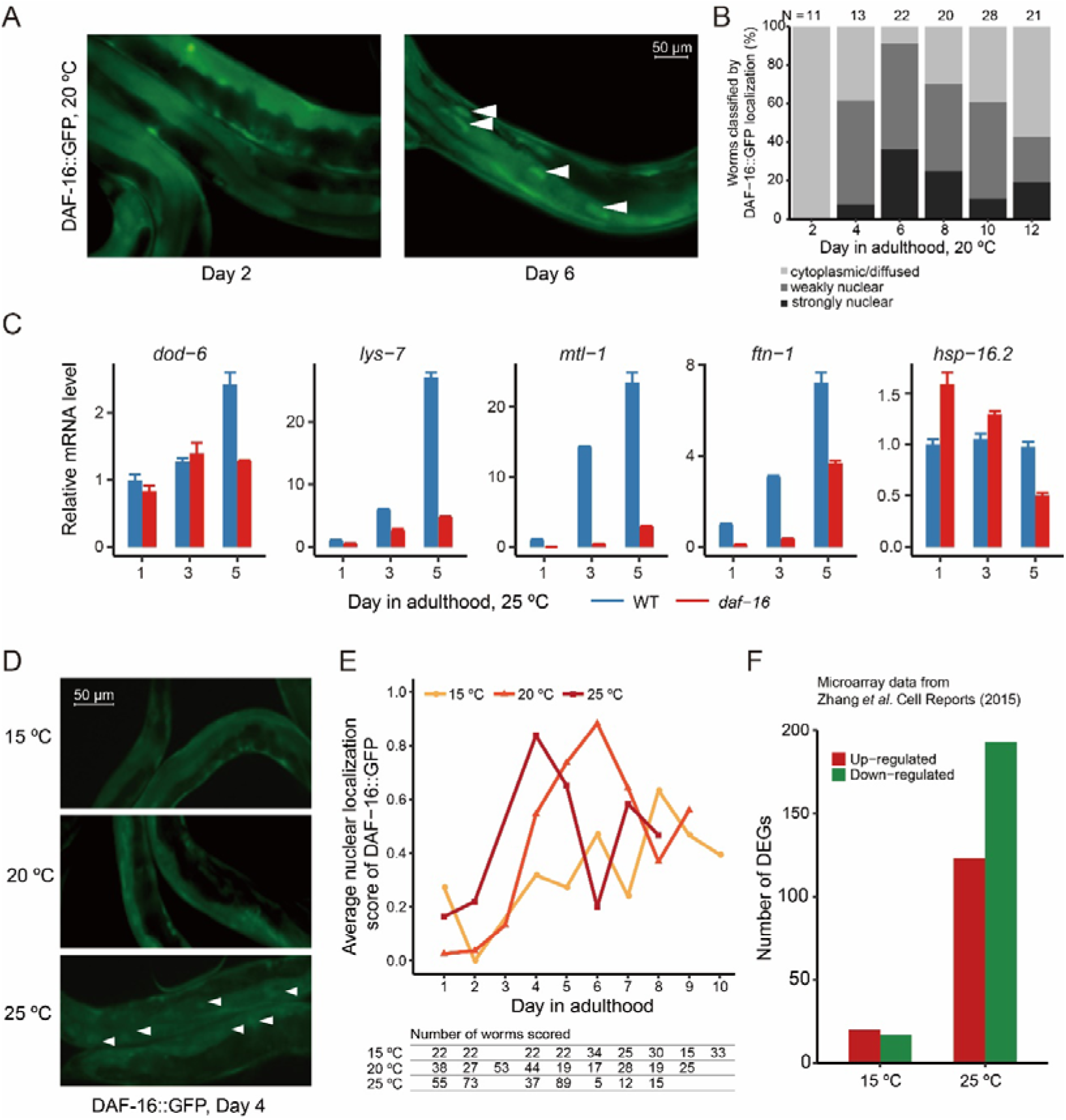
Activation of DAF-16 during the normal aging process. (A and B) Age-dependent nuclear accumulation of DAF-16::GFP in wild-type *C. elegans*. Representative images (A) and the quantitation result (B) are shown. More images are shown in Fig. S1. (C) Quantitative RT-PCR analysis of known DAF-16 target genes, expressed as mean ± standard error. *n* = 3 (D and E) An increase of culture temperature accelerates DAF-16 nuclear accumulation. Representative images and the quantitation result are shown in (D) and (E), respectively. The nuclear localization score is described in Materials and Methods. (F) Re-analysis of published microarray data (B. Zhang et al., 2015). Plotted are the numbers of DEGs between wild-type and *daf-16*(*mgDf47*) worms on adult day 3 at either 15 °C or 25 °C.

### Effect of DAF-16 activation on age-associated gene expression changes

To fully characterize the phenomenon of DAF-16 activation during aging, we performed mRNA-seq analysis of WT and *daf-16(mu86)* null mutant worms at high temporal resolution. Samples were collected every 24 hours from adult 1 through day 7 at 25 °C. The *fer-15(b26ts)* allele, which causes temperature sensitive sterility but does not affect lifespan, was used in the background to avoid sample contamination by progeny (Sijen et al., 2001). For WT worms, using adult day 1 as reference, we found that the number of DEGs increased over the next six days (Fig. 2A). For *daf-16(mu86)* null animals, the increase of DEGs was much more abrupt (Fig. 2B). To find out among the genes showing age-associated expression changes which ones were dependent on *daf-16*, we compared the transcriptomes of WT and *daf-16(null)* worms from day 1 through day 7 (Fig. 2C). As shown, there are few DEGs between WT and *daf-16(null)* on day 1, verifying that DAF-16 activity in WT worms is inhibited, which is consistent with the cytoplasmic distribution pattern of DAF-16::GFP on day 1. We defined age- and *daf-16-*dependent DEGs for each day from day 2 to day 7 as those in Regions A and B in the Venn diagram illustration in Fig. 2D. Genes in Region A are those whose expression levels increased (or decreased) significantly on day *x* (*x* > 1) compared to day 1 only in the WT, and at the same time their expression levels were significantly higher (or lower) in the WT than in the *daf-16(null)* on day *x*. For genes in Region B, their expression levels increased (or decreased) significantly on day *x* relative to day 1 in both WT and *daf-16(null)* animals, but the increase (or decrease) in the WT was significantly higher than that in the *daf-16(null)* mutant. The number of age- and *daf-16*-dependent DEGs defined as such increased from a total of 43 to a total of 802 on day 7 (Fig. 2D,E), and the daily percentage of them among age-dependent DEGs followed a similar trend rising from 1.2% on day 2 and peaking at 12.3% on day 6 (Fig. 2F). Besides the DEGs in Regions A and B, those in C and D are also *daf-16* dependent in some aspects. Comparing the transcriptome of WT on day *x* relative to either WT on day 1 or the *daf-16* mutant on day *x*, we found that the Spearman’s correlation increased steadily from 0.03 on day 2 to 0.46 on day 7, suggesting that the *daf-16* gene activity contributes more and more to age-associated transcriptome remodeling (Fig. S2). These mRNA-seq results fully supported our earlier observation (Fig. 1) of age-associated DAF-16 activation, and captured its effect on the dynamics of gene expression at 24-hour resolution for the whole genome.

**Figure 2.**
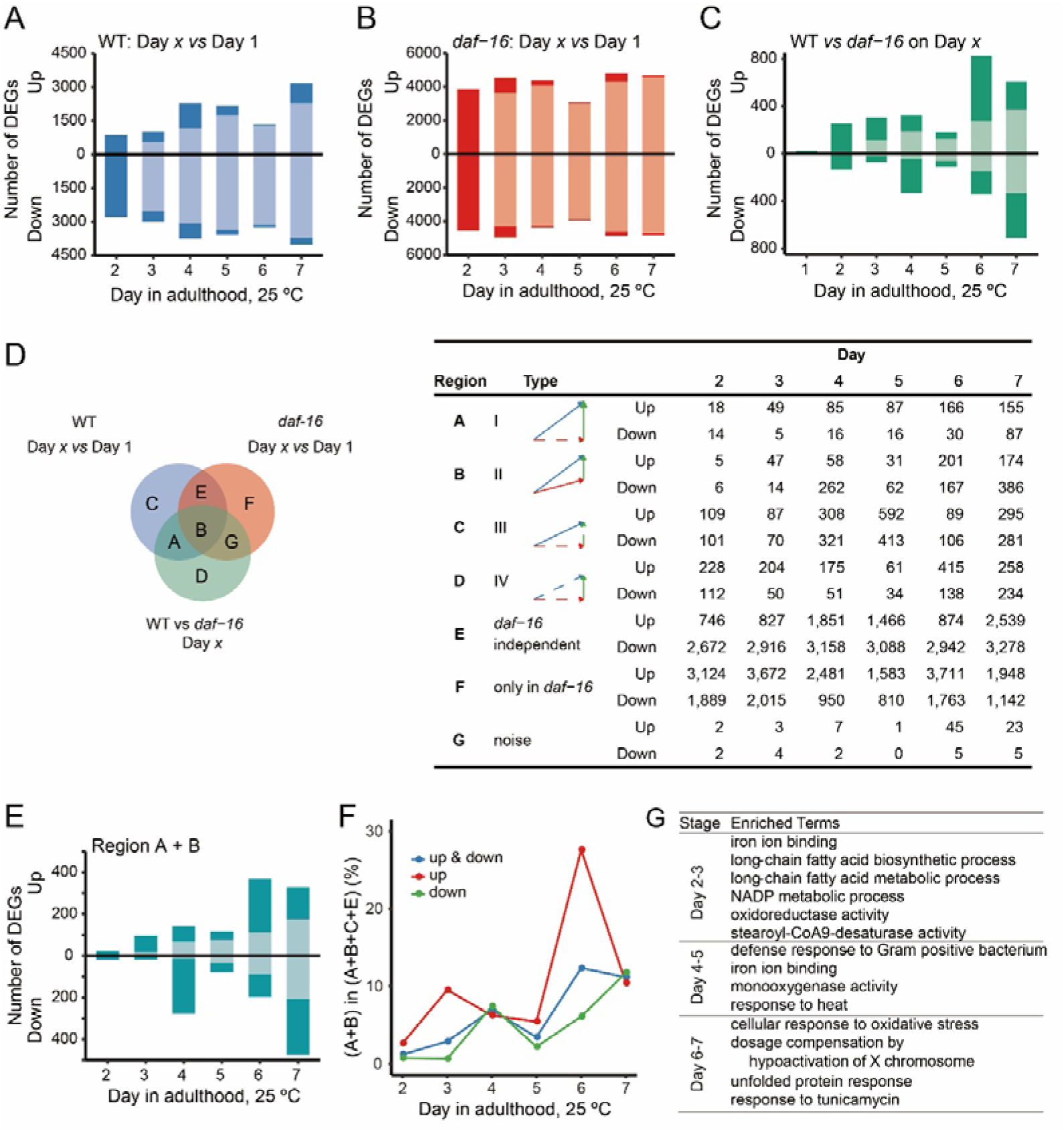
Transcriptome analysis of WT and *daf-16(null)* adults from day 1 through day 7. (A) Number of genes that are up- or down-regulated in WT on day *x* relative to day 1. Darker shades, DEGs detected for the first time; lighter shades, DEGs detected at a previous time point. (B) Number of genes that are up- or down-regulated in *daf-16(mu86)* worms on day *x* relative to day 1. (C) Number of genes that are up- or down-regulated in WT relative to the *daf-16(mu86)* mutant on each day from day 1 through day 7. (D) A mock Venn diagram to illustrate the subsets of DEGs according how they are affected by age and by *daf-16(mu86)* (left) and the number of DEGs of each subset on each day (right). A blue arrow denotes a change between Day 1 and Day *x* for WT, and a red one for the *daf-16* mutant. A green arrow denotes the difference between WT and the *daf-16* mutant on Day *x*. Statistically significant differences are indicated by solid arrows, otherwise by dashed arrows. (E) Number of age- and DAF-16-dependent DEGs (Region A+B) on Day *x* relative to Day 1. (F) Percentage of age- and DAF-16-dependent DEGs at the indicated day of age for WT *C. elegans*. This is calculated as (Region A+B) / (Region A+B+C+E) x 100%. (G) Enriched GO terms among the up-regulated genes in Region A+B. Corrected *P*-value < 0.05, fold enrichment > 2.

The mRNA-seq analysis also revealed that DAF-16 plays a role in stabilizing the transcriptome against perturbations brought about by aging. Without *daf-16*, adding only one day of age on top of day 1 was enough to change the expression of 8446 genes, whereas in the presence of *daf-16*, it took five more days to reach 7195 DEGs (Fig 2. A,B). On each day after day 1, thousands of DEGs were found only in the *daf-16* mutant (Region F), greatly outnumbered the DEGs found only in the wild type (Regions A+C) (Fig. 2D).

For nearly all the DEGs uniquely found in the *daf-16* mutant (Region F), their expression levels also increased or decreased in the WT after day 1, but not statistically significant enough to be classified as DEGs in the WT. For all genes, the fold change of mRNA reads on day *x* relative to day 1 varied to a greater extent in the *daf-16* mutant than in the WT (Fig. S3).

In conclusion, the *daf-16* gene activity seems to fulfill the function of a “capacitor” for thousands of genes to resist age-associated transcriptional alterations. This function was not known before in the normal aging process and it contrasts intriguingly with the regulator function of *daf-16*, which activates or represses the expression of a few hundred genes after day 1 (Regions A+B). The regulator function of *daf-16* directly shapes the age-dependent expression profile of WT but the capacitor function is hidden from view until *daf-16* is deleted.

### The Age-dependent DAF-16 targets are different from the IIS-dependent DAF-16 targets

One immediate question about age-associated activation of DAF-16 is whether this is caused by a gradual reduction of IIS. Arguing against this idea, the DAF-2 protein level has been found to increases with age (Tawo et al., 2017). As increased DAF-2 abundance does not necessarily lead to increased IIS and reduced DAF-16 activity, we resorted to examining the expression of the downstream targets of IIS in the normal aging process. The transcriptional targets of DAF-16 in response to reduced insulin signaling have been analyzed comprehensively by Tepper *et al*., leading to the discovery of 1663 genes that are up-regulated (Class I) and 1733 genes that are down-regulated (Class II) by DAF-16 (Tepper et al., 2013). If IIS increases with age, then expression of Class I genes should decrease and that of Class II genes should increase with age. By examining the collective temporal expression profiles of the most highly ranked class I and class II genes, we found that as WT worms become old, the top 100 class I genes tend to be up-regulated, and their class II counterparts are down-regulated (Fig. 3A). Therefore, the increase of DAF-2 protein during aging is not accompanied by inhibition of DAF-16. Further, since the age-dependent induction of the top 100 Class I genes is dependent on *daf-16*, while repression of Class II genes is independent of *daf-16* (Fig. 3A), we performed a *k*-means clustering analysis of all Class I genes (Fig. 3B). Although the 424 genes in clusters 1 and 2 support the idea of reduced IIS during aging as they show a trend of age-dependent up-regulation, the other four clusters, which consist of nearly 1000 genes, do not. Collectively, these results argue that DAF-16 activation during aging cannot be explained by either activation or inactivation of the IIS pathway.

**Figure 3.**
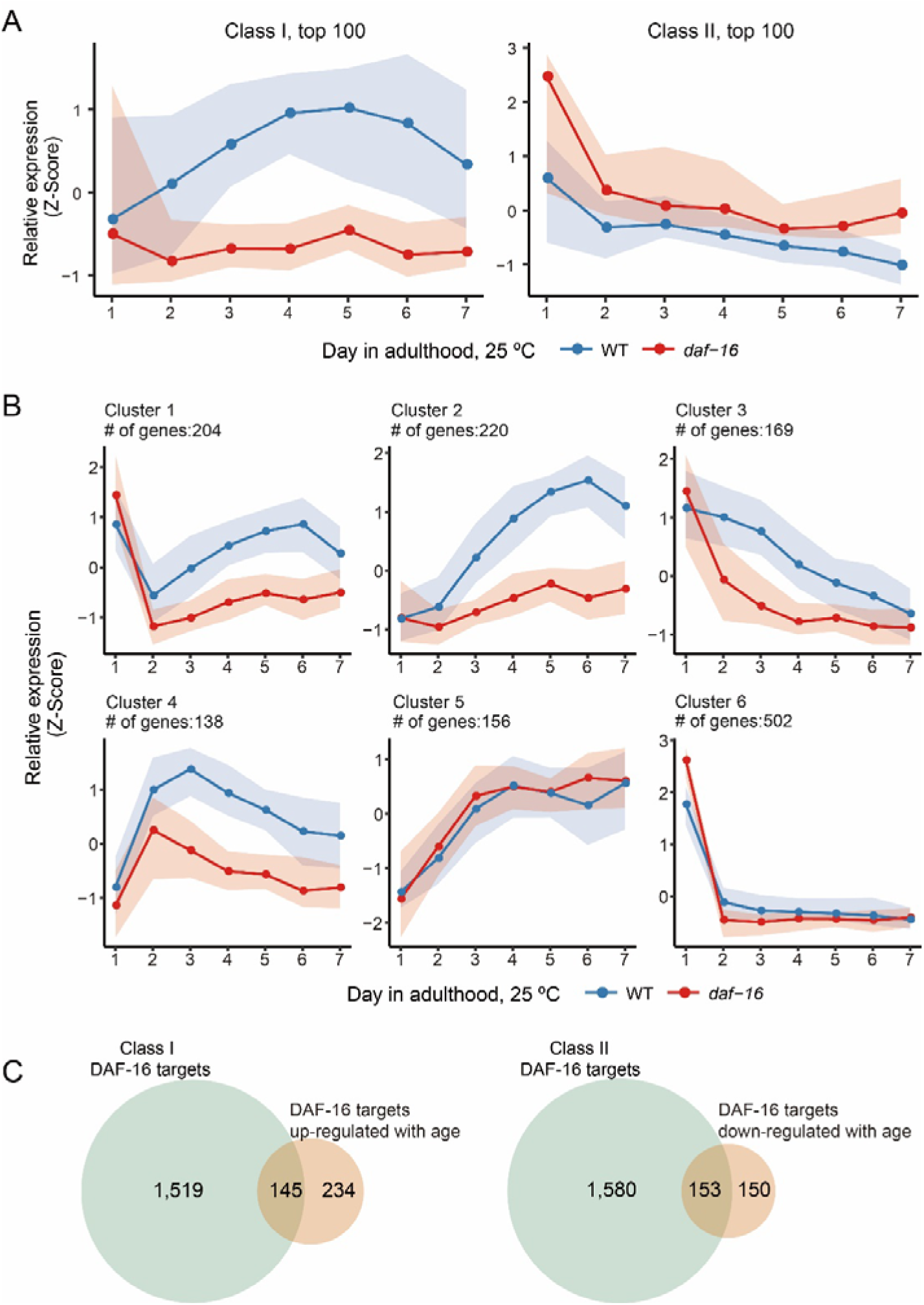
Genes regulated by DAF-16 during aging are different from the classic insulin signaling targets. (A) Relative expression (Z-Score) of the most prominent IIS targets in the first week of adulthood. Left panel: top 100 Class I genes; right panel: top 100 Class II genes. (B) *k*-means clustering of the temporal expression profiles of Class I genes. For both (A) and (B), the lines and the shade boundaries indicate the 50^th^, 20^th^ and 80^th^ percentile. The expression levels of each gene across all samples are centered to the mean and scaled to the variance. (C) Overlap of the classic IIS targets and the genes regulated by DAF-16 during aging.

To compare age-dependent DAF-16 targets with IIS-dependent DAF-16 targets, we defined age-dependent DAF-16 target genes as those whose expression levels were significantly higher or lower in the WT than in the *daf-16(mu86)* mutant on both day 6 and day 7, and on both days the fold change of WT vs *daf-16(mu86)* must be greater than that on day 1. As expected, the overlap between class I and 379 DAF-16 targets up-regulated by age is not extensive, neither is the overlap between class II genes and 303 DAF-16 targets down-regulated by age (Fig. 3C). All combined, less than 50% of the age-dependent DAF-16 targets were previously identified as DAF-16 targets in response to reduced IIS.

### Activation of DAF-16 in response to age-associated proteostasis collapse

Next, we tried to find out what may be the cause of DAF-16 activation in the normal aging process. Besides IIS, DAF-16 can be regulated by other signals in the cellular environment, including various stress signals such as heat, oxidative stress, and starvation (Henderson & Johnson, 2001). Since different types of stress are typically coupled together during the aging process, discerning cause and consequence remains the most challenging in aging research. In this study, we focused on the collapse of proteostasis as it is shown to be an early and profound aging phenotype in *C. elegans* and is proposed to be a driving force of aging (Ben-Zvi, Miller, & Morimoto, 2009). The collapse of proteostasis induces the expression of various heat-shock proteins (HSPs), which help refold damaged proteins. From our mRNA-seq data, we found evidence of *daf-16* dependent induction of multiple small HSPs a few days into the adulthood (Fig. 4A).

**Figure 4.**
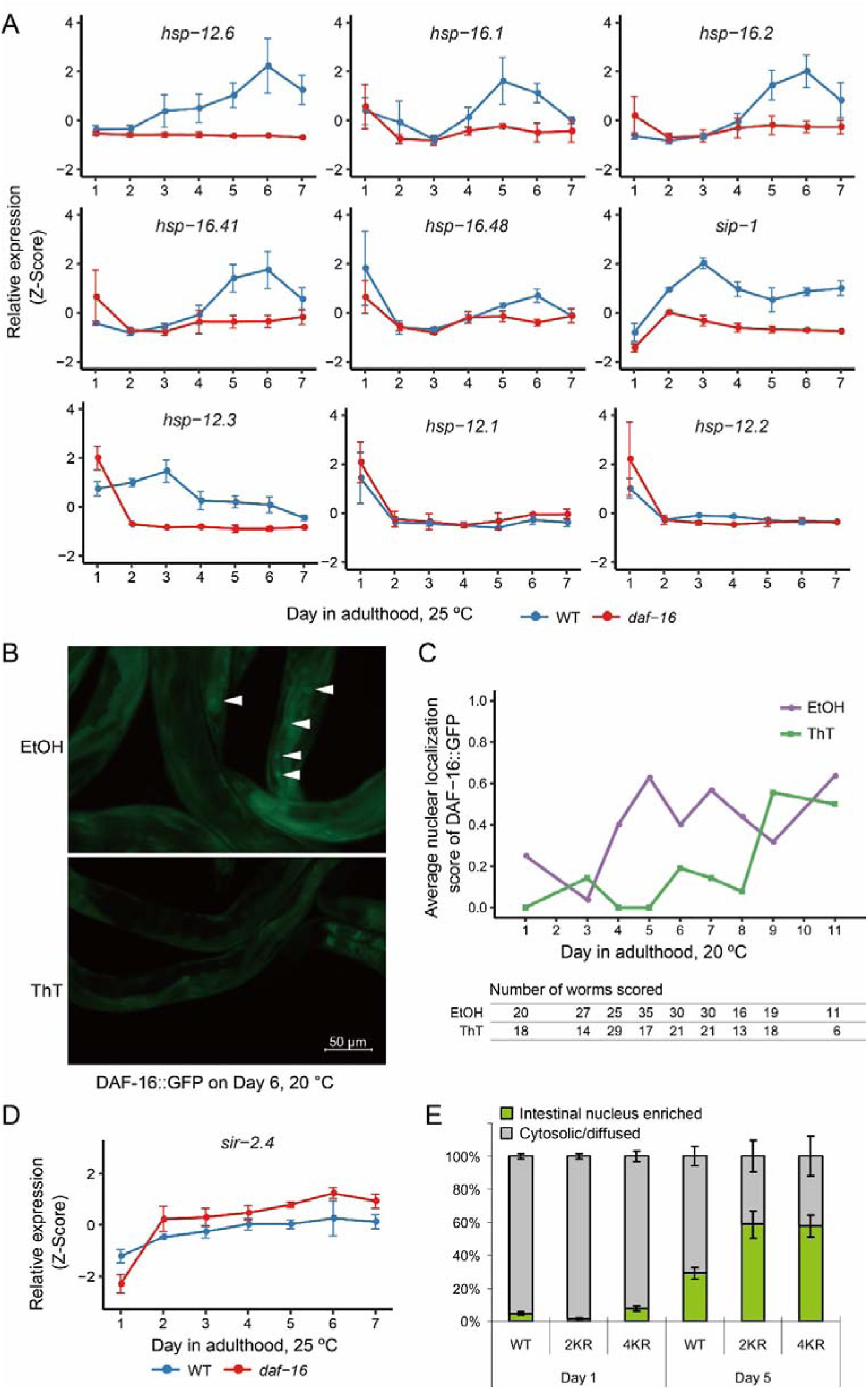
Activation of DAF-16 during aging is connected to the loss of proteostasis. (A) Temporal expression profiles of the genes encoding small heat shock proteins. (B and C) Thioflavin T (ThT) attenuated age-dependent DAF-16 nuclear localization. Representative images (B) and the quantitation result (C) are shown. (D) The temporal expression profile of *sir-2.4*. (E) DAF-16 K to R mutations, which abolish acetylation and sensitize DAF-16 for nuclear translocation, enhanced age-dependent nuclear accumulation of DAF-16::GFP

Additionally, gene ontology (GO) analysis of age- and DAF-16-dependent DEGs have suggested that among the up-regulated DEGs identified from day 4 to day 7, a number of stress response terms are significantly enriched (Fig. 2G). For example, response to heat and defense response to Gram-positive bacterium are enriched most prominently on day 4 and day 5; unfolded protein response, cellular response to oxidative stress, and response to tunicamycin (ER stress) (Olden, Pratt, Jaworski, & Yamada, 1979) are enriched on day 6 and day 7.

This suggests that the collapse of proteostasis is a plausible cause of DAF-16 activation in older worms particularly on day 6 and day 7. To test this hypothesis, we treated *C. elegans* with Thioflavin T (ThT), a compound that improves proteostasis and extends lifespan by binding to and stabilizing amyloids (Alavez, Vantipalli, Zucker, Klang, & Lithgow, 2011). Indeed, ThT treatment delayed nuclear accumulation of DAF-16::GFP by five days at 20 °C (Fig. 4B,C), suggesting that the collapse of proteostasis is a cause for DAF-16 activation in older worms, although it may not be the only one.

SIR-2.4, the *C. elegans* homolog of mammalian SIRT6 and SIRT7, promotes stress-induced DAF-16 nuclear localization by removing the acetyl modification groups from several lysine residues of DAF-16 (Chiang, Tishkoff, et al., 2012). Our mRNA-seq experiments showed that the expression of *sir-2.4* was elevated in older worms. Induction of *sir-2.4* is in favor of promoting DAF-16 nuclear localization. Consistently, 2KR and 4KR mutations, both of which preclude CBP-1-dependent acetylation and thus sensitize DAF-16, enhanced age-dependent nuclear localization of DAF-16::GFP (Fig. 4E). To sum up, the evidence above strongly suggests a mechanism of DAF-16 activation in response to age-associated cellular stress.

### Additional transcription factors cooperate with DAF-16 to shape the aging transcriptome

Lastly, we asked what additional transcription factors besides DAF-16 may take part in age-associated transcriptome remodeling. The mRNA-seq analysis uncovered thousands of DAF-16 independent DEGs from day 2 to day 7, compared with day 1 (Region E, Fig. 2D). Also, there are DEGs that are only partially dependent on DAF-16 (Region B, Fig. 2D). Expression of *daf-16* itself increases with age (Fig. 5A), which is likely to be the result of transcriptional activation by other transcription factors, because *daf-16* is not a class I gene, i.e., not a target of positive regulation by its own gene product (Tepper et al., 2013). Thus, there must be additional TFs that govern age-dependent transcription. We examined the temporal expression profiles of seven additional TFs that have been shown to play a role in lifespan regulation (Fig. 5A). In both WT and *daf-16* mutant worms, expression of *hsf-1* and *skn-1* increased with age, while expression of *elt-2* decreased with age. In contrast, the age dependent increase of *pqm-1* expression was dependent on *daf-16*, and so was that of *hlh-30*. The temporal expression profiles of *pha-4* and *dve-1* were more complex. We wondered whether *hsf-1* and *skn-1* cooperate with *daf-16* during aging, because they are critical for stress response (An & Blackwell, 2003; Hsu, Murphy, & Kenyon, 2003) and their expression levels increase with age. Using the transcriptional reporter strains of *dod-3*, *mtl-1* and *hsp-16.2*, we found that *hsf-1* is required for the enhanced expression of *hsp-16.2p::nCherry* in mid-aged worms, but not for that of *dod-3p::gfp* and *mtl-1p::bfp* (Fig. 5B,C). *skn-1* RNAi had no effect on all three. Six well characterized *skn-1* target genes *gst-4/5/7/10/13/38*) (Oliveira et al., 2009; Tullet et al., 2008) are generally down-regulated during aging, indicating a loss of SKN-1 transcriptional activity despite the increase of *skn-1* mRNA, consistent with a previous finding (Ewald, Landis, Porter Abate, Murphy, & Blackwell, 2015). We also examined the effect of *elt-2* RNAi, as *elt-2* has been reported to regulate insulin signaling targets and is responsible for age-dependent down-regulation of several hundred genes (Mann et al., 2016; P. Zhang, Judy, Lee, & Kenyon, 2013). Indeed, *elt-2* was required for the induction of *dod-3p::gfp* and *mtl-1p::bfp* expression in mid-aged worms (Fig. 5B,C). Therefore, at least two TFs, ELT-2 and HSF-1, cooperate with DAF-16 to regulate different sets of target genes during aging.

**Figure 5.**
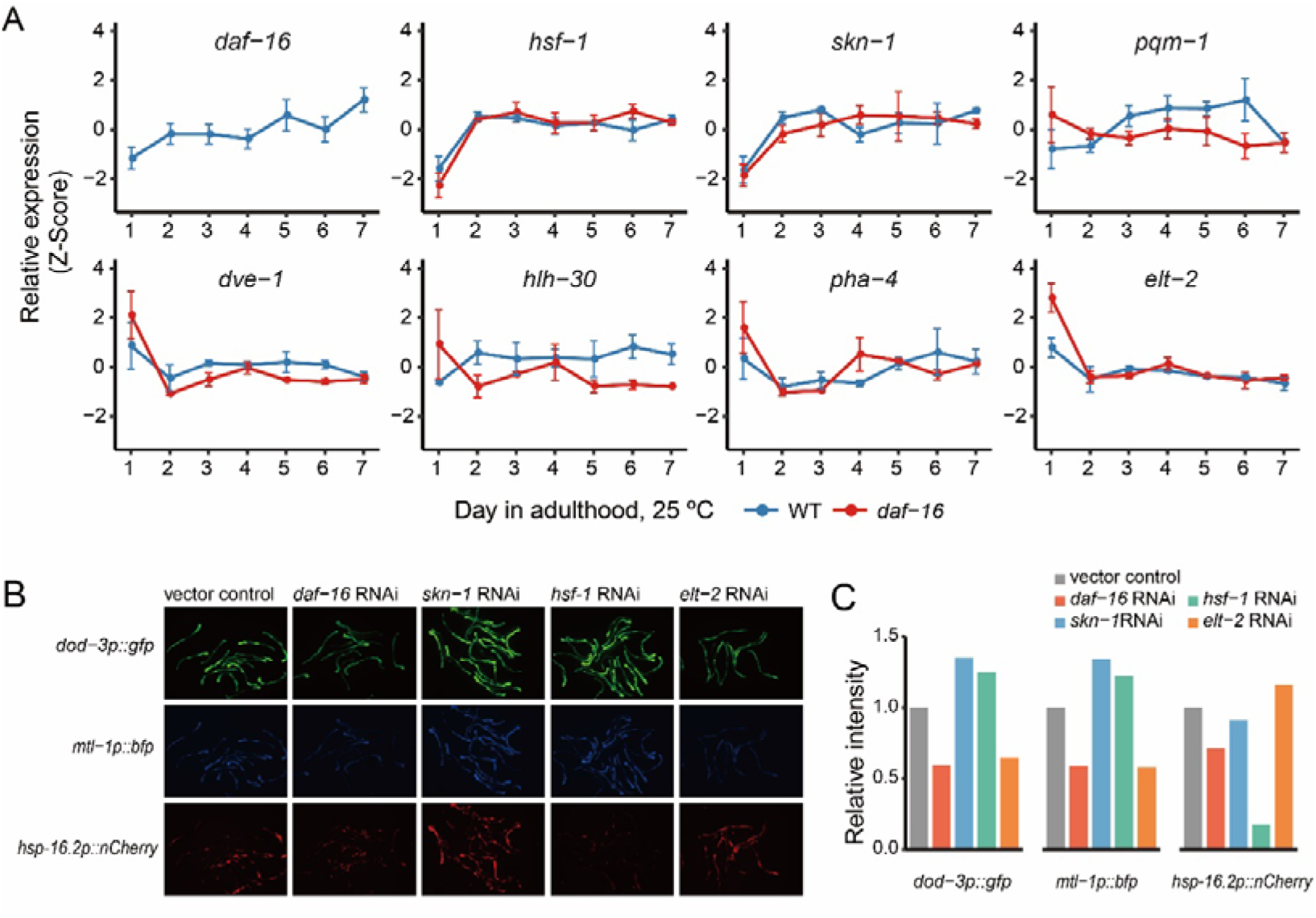
Additional Transcription factors cooperate with DAF-16 to shape age-dependent gene expression changes. (A) Expression profiles of *daf-16* and seven additional transcription factors as seen in our mRNA-seq analysis. (B and C) Knocking down some of the transcription factors by RNAi affected the expression of *dod-3*, *mtl-1*, and *hsp-16.2* on adult day 5 as indicated by fluorescent reporters. Representative images (B) and the quantitation result (C) are shown.

## Discussion

### DAF-16 activation in mid-aged worms indicates that aging is not entirely passive or haphazard

Aging is intriguing. It seems inevitable for animals that age, even if they are given the best living conditions possible. One school of evolutionary theories including the widely known antagonistic pleiotropy theory attribute the origin of aging to a declining force of selection on individuals after reproduction begins (Trindade et al., 2013; Williams, 1957). The weakening or lacking of purifying selection eventually takes out proper regulation of a living system and let mutations that have detrimental effects late in life accumulate in the genome. Based on these theories, one might expect the aging process to unfold in a passive and haphazard way. However, our high-resolution transcriptome analysis of *C. elegans* adults at 25 °C from day 1 through day 7 has depicted a different view. Activation of DAF-16 is detectable by mRNA-seq starting from day 2 and becomes well pronounced after day 5 (Fig. 1E, Fig. 2C and Fig. S2) when reproduction has ended and aging undoubtedly kicks in. Activation of DAF-16 turns on the expression of many stress response genes as is evident from the significantly enriched GO terms (Fig. 2G) such as defense response to Gram-positive bacterium, response to heat, response to oxidative stress, unfolded protein response, and response to tunicamycin (ER stress). Thus, DAF-16 is activated in the normal aging process for protection and repair, i.e., anti-aging. Previous studies have identified transcription factors such as ELT-2 and ELT-3 whose deactivation during aging drives the transcriptional network into disorder (Budovskaya et al., 2008; Mann et al., 2016). Here, by showing DAF-16 activation after adult day 1, we provide evidence for active gene expression remodeling during normal aging. This shows that the aging process is not entirely passive. Also, based on the GO terms of the DAF-16 targets that are turned on from day 2 to day 7, this period may be divided into three phases. The first one, of days 2-3, is significantly enriched for GO terms indicating metabolic activities; the third one, of days 6-7, is significantly enriched for GO terms of stress response; and in between is the transition phase (Fig. 2G). When aging starts is a deep question—it is difficult to draw a hard line. Nevertheless, at 25 °C reproduction finishes at the end of day 3, so one can argue that by this time, the force of natural selection on individual *C. elegans* has disappeared, and aging has the stage all to itself. If day 4 is the start line, then aging starts with the transition phase followed by a stress response phase, and probably others that have not been characterized. This shows that the aging process is not entirely haphazard, either, it proceeds in an orderly fashion at least in the beginning.

Taken together, our surprising finding of DAF-16 activation during normal aging indicates that aging is not a passive and haphazard process as often perceived before, as least not entirely. There is active anti-aging regulation and there is order among disorder.

### Distinct transcriptional output of DAF-16 during aging

By defining nearly 700 age-dependent DAF-16 targets (Fig. 3C), we were surprised to find that more than half of them were not identified before as DAF-16 targets downstream of IIS. By KEGG analysis of the IIS-regulated and age-regulated DAF-16 target genes, we find that drug metabolism, porphyrin and chlorophyll metabolism, and glycerolphopholipid metabolism are specifically enriched among the targets that are down-regulated by DAF-16 during aging (Fig. S4). Of interest, genes involved in mTOR signaling or ErbB signaling are specifically enriched among the targets up-regulated by DAF-16 during aging (Fig. S4). In mammalian cells IIS regulates, and is regulated by, the mTOR pathway (Laplante & Sabatini, 2012), and both pathways critically regulate metabolism and aging. However, the connection between these two signaling pathways in *C. elegans* is unknown. No clear evidence has been found to place the *C. elegans* mTOR pathway downstream of IIS. Our transcriptome analysis now finds mTOR signaling downstream of DAF-16 during aging. It awaits further investigation to uncover the underlying mechanism.

### DAF-16 activation in response to the loss of proteostasis

DAF-16 is known to integrate multiple genetic and environmental stimuli (Kenyon 2010), and this study uncovers aging as another one. DAF-16 activation during aging produces a set of transcriptional output different from that by reduction of IIS. *C. elegans* and human have a long post-reproductive lifespan. In both systems, post-mitotic cells live for a long time, either in absolute timing units (e.g. neurons and cardiomyocytes in human) or relative to the length of the reproductive period (all somatic cells in *C. elegans*). Without the ability to dilute damaged proteins by cell division, little is known about how these cells handle the loss of proteostasis, a hallmark of aging (López-Otín, Blasco, Partridge, Serrano, & Kroemer, 2013). In human, repressor element 1-silencing transcription factor (REST), which is often lost in Alzheimer’s patients, is activated in healthy seniors to induce the expression of stress response genes and to repress the genes that promote cell death and Alzheimer’s (Lu et al., 2014). This suggests that cellular stress response plays an active role in healthy aging. Consistently, this study finds that the loss of proteostasis is a cause of DAF-16 activation during normal aging. Modulating proteostasis by temperature or amyloid binding compounds changes the timing of DAF-16 nuclear accumulation. It has been shown that *hsf-1* mediated heat shock response is repressed from early adulthood (Labbadia & Morimoto, 2015), so the stress response mediated by DAF-16 in mid-aged worms, which includes the induction of small HSPs, implicates an arms race between cellular protection mechanisms and aging.

### Role of DAF-16 in wild-type *C. elegans*

The *daf-16(null)* mutants live slightly shorter than WT on standard culture plates at 20 °C (Lin, Hsin, Libina, & Kenyon, 2001). In comparison, loss of *daf-16* produces a dramatic effect in the *daf-2* mutant background as *daf-16(null)* completely abolishes the exceptional longevity of *daf-2* mutant worms (Kenyon et al., 1993). Besides, unlike *daf-2(null)* mutants, which are lethal, *daf-16(null)* mutants are perfectly viable and fertile. This makes one wonder what is the purpose of *daf-16* in WT *C. elegans*. Here we show that although DAF-16 is largely inactive on adult day 1, judging by the fact that there is little difference between WT and *daf-16(null)* at this point (Fig. 2C), DAF-16 acts as a “capacitor” to stabilize the transcriptome during aging (Compare Fig. 2A and Fig. 2B).

### Timing of DAF-16 activation for longevity

Previously, an elegant genetic study has demonstrated that for the longevity phenotype induced by reduction of IIS, activation of *daf-16* in early adulthood is the most critical (Dillin, Crawford, & Kenyon, 2002). In this study, we find that *daf-16* is activated in mid-aged WT animals. This is consistent with the timing requirement of lifespan regulation by IIS, and explains why late activation of DAF-16 by reducing IIS is less effective—because DAF-16 is already activated.

## Experimental Procedures

### C. elegans Strains

Strains used in this work include N2, CF1038 *daf-16(mu86) I*, MQD54 *hqIs9*[*hqIs9*[*daf-16p::DAF-16::6xHis::GFP, pRF4*], DH26 *fer-15(b26ts) II*, MQD1319 *daf-16(m86) I; fer-15(b26ts) II* and MQD1586 *hqEx476*[*hsp-16.2p::nCherry; dod-3p::gfp; mtl-1p::bfp, unc-119(+)*]; *unc-119(ed3) III* EQ1137 *daf-16(mu86) I; iqIs79*[*pAH123(daf-16p::daf-16::gfp^wt^, pRF4*]; EQ1079 *daf-16(mu86) I; iqIs83*[*pAH144(daf-16p::daf-16^4KR, K246+251+373+377R^)::gfp, pRF4*]; EQ1070 *daf-16(mu86) I; iqIs83*[*pAH144(daf-16p::daf-16^4KR, K246+251+373+377R^)::gfp, pRF4*]

To generate transgenic animals, a plasmid DNA mix was microinjected into the gonad using standard method. The plasmid DNA mix consisted of 30 ng/µl of the indicated plasmid and 80 ng/µl of the pRF4 or 10 ng/µl of the pCFJ151. The UV irradiation was used to integrate the extra-chromosomal DNA into the genome. The integrated strains were further back-crossed 6X to CF1037 *daf-16(mu86) I*. MQD1319 were generated from CF1038 *daf-16(mu86) I* and DH26 *fer-15(b26ts) II* and backcrossed to DH26 for four times.

### DAF-16 nuclear localization

For the effect of the culture temperature on DAF-16::GFP localization, worms were cultured on standard Nematode Growth Medium (NGM) plates. For ThT treatment, L4 larvae grown at 20 °C were transferred to NGM plates containing 50 μM ThT (SIGMA, St. Louis, MO, USA) dissolved in ethanol. NGM plates with equal amount of ethanol served as the control. As soon as worms were removed from incubation, they were mounted on slides and imaged immediately. The images were taken using a Zeiss Axio Imager M1 microscope at 400-fold magnification. For the ThT experiments, DAF-16::GFP fluorescent images were taken using the YFP channel to avoid interference of ThT fluorescence. All other fluorescent image of DAF-16::GFP nuclear were taken using the GFP channel.

For the effect of the culture temperature and ThT on DAF-16::GFP localization, each worm was given a score: cytoplasmic, 0; weakly nuclear, 1; strongly nuclear, 2.

For the analysis of nuclear localization in KR mutant worms, GFP localization was then analyzed using an Olympus BX61 fluorescent microscope on adult day 1 and day 5. An animal was scored as having nuclear GFP if more than one intestinal nucleus contained enriched GFP signal. The quantification of DAF-16::GFP localization was described previously in (Chiang, Ching, Lee, Mousigian, & Hsu, 2012).

### Quantitative real-time PCR (qRT-PCR)

Total RNAs were extracted from WT or *daf-16(mu86)* worms of the indicated age using TRIZOL (INVITROGEN, Grand Island, NY, USA), followed by the removal of contaminant DNA using DNase I. cDNAs were synthesized from total RNA templates using a reverse transcription kit (TAKARA, Kusatsu, Shiga, Japan). qPCR was carried out on an ABI 7500 Fast real-time PCR system using a TAKARA real-time PCR kit (SYBR Premix Ex TaqTM II). *pmp-3* was used as the internal control. The qPCR primers were:

*mtl-1* [TGAGGAGGCCAGTGAGAAAAA]/[GCTCTGCACAATGACAGTTTGC];

*ftn-1* [TGACGCGCACTTGACAAATTA]/[TGTAGCGAGCAAATTCATTGATC];

*dod-6* [CTCAAGACCGTCGCCCTCTA]/[TCAGCATCAGCGCAAGCA];

*lys-7* [CATTCGGCATCAGTCAAGGTT]/[GCAGGCTCCGCAATGACTT];

*hsp-16.2* [TACGCTATCAATCCAAGGAGAAC]/[GAAGCAACTGCACCAACATC];

*pmp-3* [GAATGGAATTGTTTCACGGAATGC]/[CTCTTCGTGAAGTTCCATAACACGATG].

### RNA sequencing and data analysis

Transcriptome analyses of *fer-15(b26ts)* and *daf-16(mu86); fer-15(b26ts*) worms were carried out in three biological replicates. Worms were grown on High Growth (HG) plates supplemented with OP50 bacteria at 25 °C and harvested on adult day 1 through day 7. RNA quality was evaluated on a Bioanalyzer 2100 instrument (Agilent, Santa Clara, CA). Sequencing libraries were prepared following the protocol of the NEBNext Ultra RNA library Prep Kit (NEB) and sequenced on an Illumina HiSeq X Ten platform in the paired-end mode (2×150 bp) through the service provided by Bionova. An average of 25M clean reads were generated for each replicate. RNA-seq reads were aligned to the *C. elegans* reference genome (ws235) using HISAT2 (v2.0.4) (Kim, Langmead, & Salzberg, 2015) with default parameters. Gene-level read counts were calculated using HTSeq (v0.6.1p1) (Anders, Pyl, & Huber, 2015) based on the Ensembl gene annotation v85. DEseq2 (v1.18.1) was used for data normalization (Love, Huber, & Anders, 2014).

Hieratical clustering of samples by Spearman’s correlation of gene expression was adapted to examine the quality of the data. Five outliers (rep 3 of *fer-15* on Day 2/3/6, rep 3 of *daf-16; fer-15* on Day 2, and rep 2 of *daf-16; fer-15* on Day 5) were removed from further analysis.

Statistical analysis of differential expression was performed using the *nbinomWaldTest* in DESeq2 package. Genes with an adjusted *p*-value < 0.05 were defined as differentially expressed genes (DEGs).

In defining age-dependent DAF-16 targets, the day-6 and day-7 DEGs were filtered by requiring an adjusted *p*-value < 0.1. Class I and II DAF-16 targets under reduced IIS are from (Tepper et al., 2013).

### Enrichment analysis

Gene Ontology (GO) and KEGG pathway databases were downloaded from the KOBAS website (Xie et al., 2011). The enrichment analysis was done using a hypergeometric test, and the *p-*values were adjusted using the BH (Benjamini & Hochberg, 1995) method.

### K-means clustering and data normalization

The normalized read counts of each gene across all samples were first transformed to Z-Score (mean centered and scaled to the variance). K-means clustering was performed using the *kmeans* function in R with the settings of 6 clusters and a maximum of 1,000 iterations.

### Fluorescence transcriptional reporter assay

All fluorescent images in Figure 5 were taken using a Zeiss Axio Imager M1 microscope at 100X magnification. Each image was taken as a whole object and the fluorescence intensity was calculated with background subtraction using image J.

### RNAi

RNAi assays were performed at 20 °C using the feeding method as previously described (Tao et al., 2013). 50 μM FudR was added into the RNAi plate to prevent the contamination of progeny. Worms were fed RNAi bacteria from adult day 1. The RNAi bacterial strain of *daf-16* was from the Ahringer RNAi library, and other RNAi bacterial strains were from the Vidal RNAi library. The *E. coli* strain HT115 transfected with L4440 (empty vector) was used as control.

### Data availability

All the FASTQ files for RNA-seq from this study will be submitted to the NCBI Sequence Read Archive (SRA; http://www.ncbi.nlm.nih.gov/sra) under accession number SRPxxxxxx.

## Acknowledgments

We thank the Ministry of Science and Technology of China (973 grant 2014CB84980001 to M.-Q. D.), the municipal government of Beijing, and Ministry of Science and Technology of Taiwan (MOST 102-2311-B-010-010-MY3 to A-L. Hsu) for funding.

## Author contributions

S.-T.L., conception and design, acquisition of data, analysis and interpretation of data, drafting and revising the article. H.-Q.Z., conception and design, analysis and interpretation of data, drafting and revising the article. P.Z. conception and design, acquisition of data. C.-Y.L., acquisition of data. Y.-P.Z., acquisition of data. A.-L.H., interpretation of data. M.-Q.D., conception and design, analysis and interpretation of data, drafting and revising the article.

## Supporting Information Listing

Figure S1. Representative images of DAF-16::GFP localization in young and old worms

Figure S2. Correlation coefficient of transcriptome comparing [Day *x*: WT to *daf-16*] *vs* [WT Day *x* to WT Day 1] reveals increased important of DAF-16 in shaping the transcriptome during aging.

Figure S3. The presence of the *daf-16* gene attenuates gene expression drift during aging.

Figure S4. KEGG enrichment of DAF-16 target genes in response to reduced IIS or those in response to aging.

Figure S5. Temporal expression profiles of six *skn-1* target genes.

