## Supplementary Materials for "Insulin Signaling-independent Activation of DAF-16 Shapes the Transcriptome during Normal Aging"

Figure S1. Representative images of DAF-16::GFP localization in young and old worms

Figure S2. Correlation coefficient of transcriptome comparing [Day *x*: WT to *daf-16* ] *vs* [WT Day *x* to WT Day 1] reveals increased important of DAF-16 in shaping the transcriptome during aging.

Figure S3. The presence of the *daf-16* gene attenuates gene expression drift during aging.

Figure S4. KEGG enrichment of DAF-16 target genes in response to reduced IIS or those in response to aging.

Figure S5. Temporal expression profiles of six *skn-1* target genes.

#### Figure S1

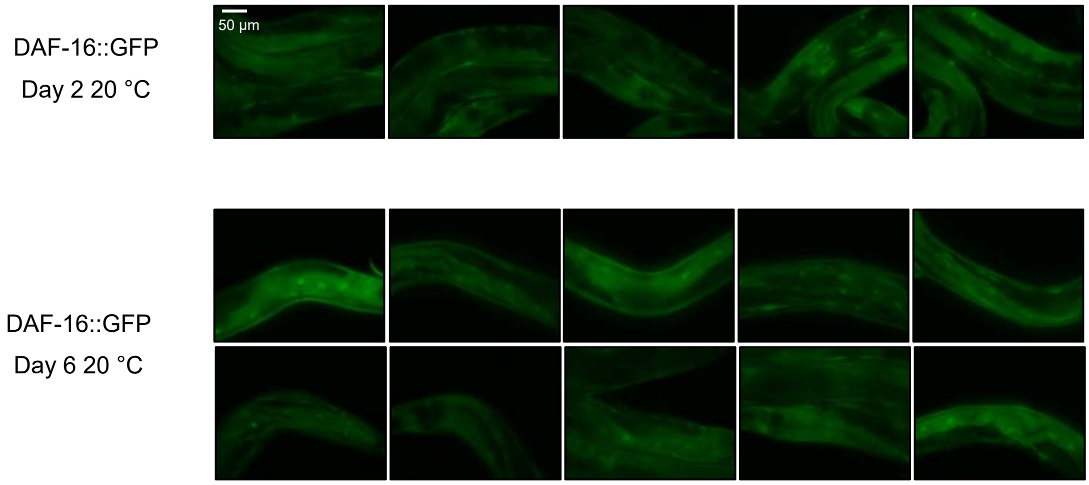


#### Figure S2

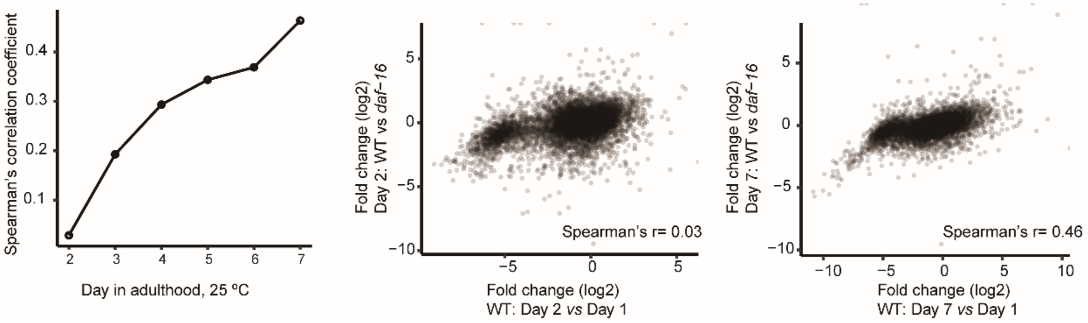

Figure S3

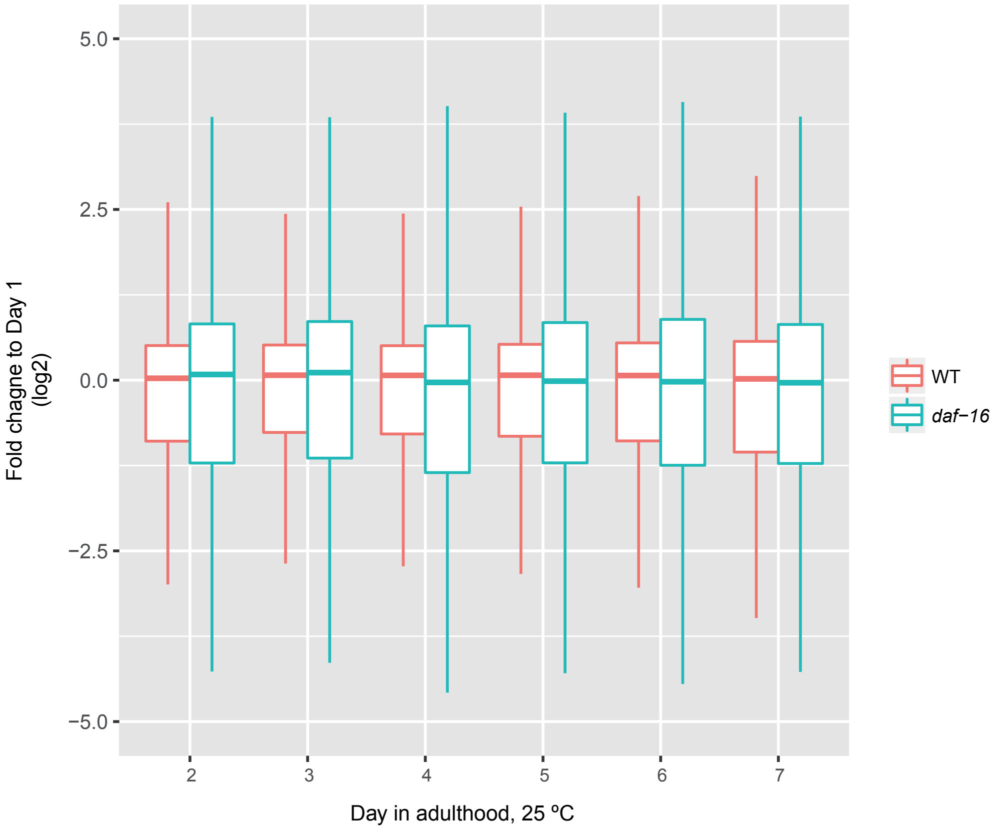
Fold chagne to Day 1
(log2)
5.0
2.5
0.0
−2.5
−5.0
2
3
4
5
6
7
Day in adulthood, 25 ºC
WT
daf−16

### Figure S4

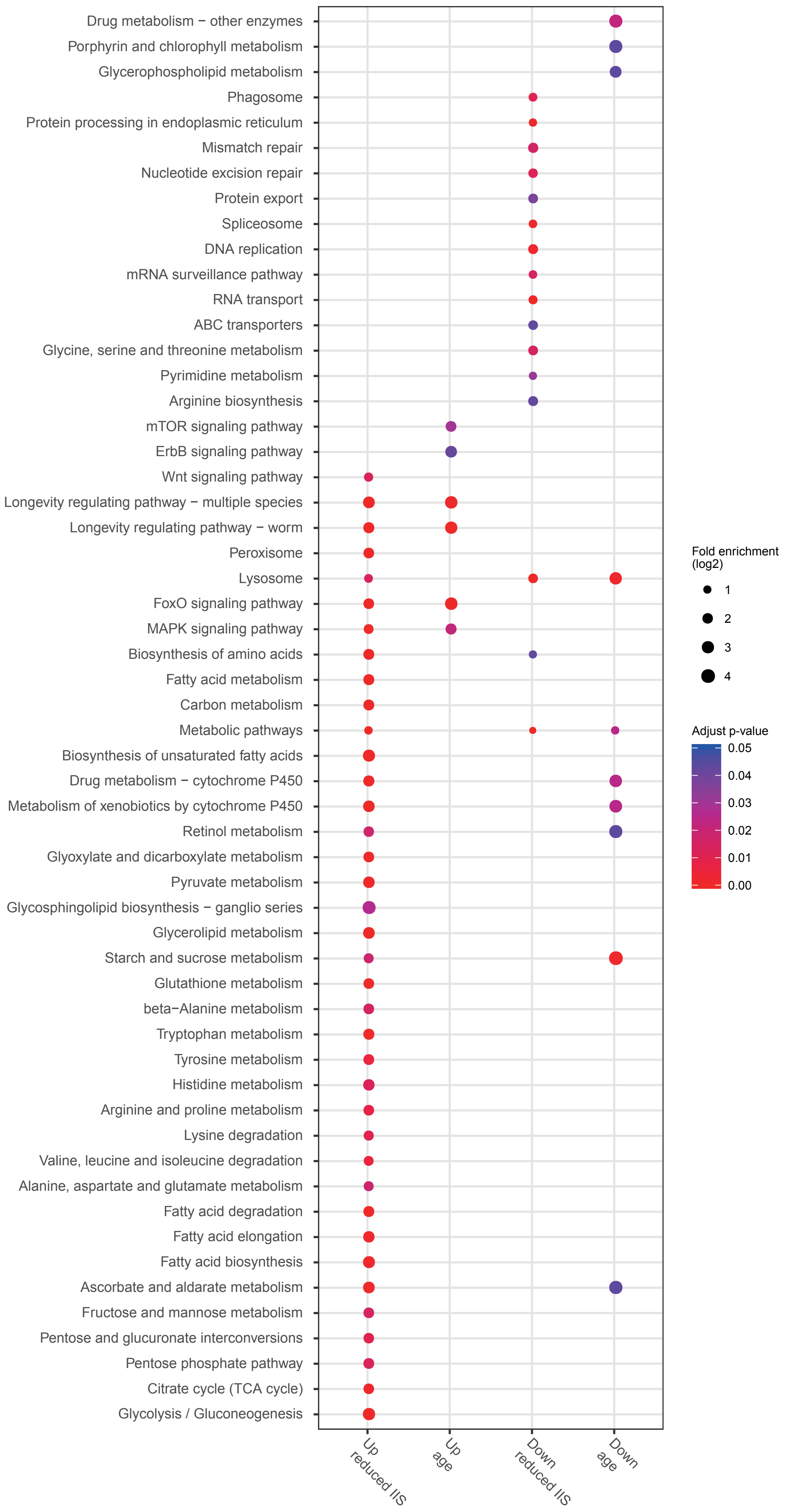
Drug metabolism − other enzymes
Porphyrin and chlorophyll metabolism
Glycerophospholipid metabolism
Phagosome
Protein processing in endoplasmic reticulum
Mismatch repair
Nucleotide excision repair
Protein export
Spliceosome
DNA replication
mRNA surveillance pathway
RNA transport
ABC transporters
Glycine, serine and threonine metabolism
Pyrimidine metabolism
Arginine biosynthesis
mTOR signaling pathway
ErbB signaling pathway
Wnt signaling pathway
Longevity regulating pathway − multiple species
Longevity regulating pathway − worm
Peroxisome
Lysosome
FoxO signaling pathway
MAPK signaling pathway
Biosynthesis of amino acids
Fatty acid metabolism
Carbon metabolism
Metabolic pathways
Biosynthesis of unsaturated fatty acids
Drug metabolism − cytochrome P450
Metabolism of xenobiotics by cytochrome P450
Retinol metabolism
Glyoxylate and dicarboxylate metabolism
Pyruvate metabolism
Glycosphingolipid biosynthesis − ganglio series
Glycerolipid metabolism
Starch and sucrose metabolism
Glutathione metabolism
beta−Alanine metabolism
Tryptophan metabolism
Tyrosine metabolism
Histidine metabolism
Arginine and proline metabolism
Lysine degradation
Valine, leucine and isoleucine degradation
Alanine, aspartate and glutamate metabolism
Fatty acid degradation
Fatty acid elongation
Fatty acid biosynthesis
Ascorbate and aldarate metabolism
Fructose and mannose metabolism
Pentose and glucuronate interconversions
Pentose phosphate pathway
Citrate cycle (TCA cycle)
Glycolysis / Gluconeogenesis
Up reduced IIS
Up age
Down reduced IIS
Down age
Fold enrichment (log2)
1
2
3
4
Adjust p-value
0.05
0.04
0.03
0.02
0.01
0.00

Figure S5

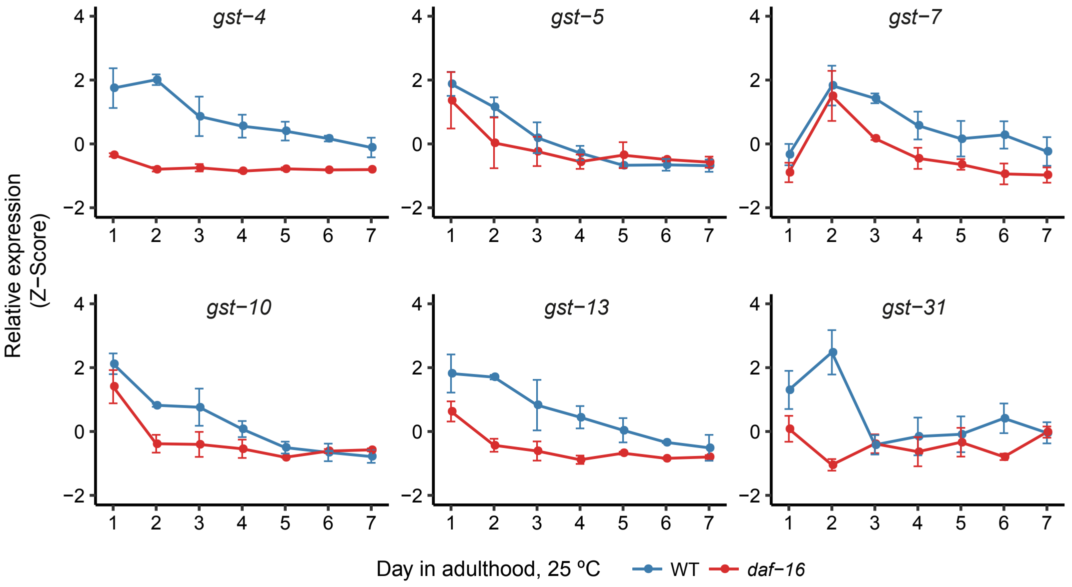

gst−4
gst−5
gst−7
gst−10
gst−13
gst−31
Relative expression
(Z−Score)
4
2
0
−2
1 2 3 4 5 6 7
Day in adulthood, 25 ºC
WT
daf−16
